# Epigenetically silenced DACT3 promotes tumor growth via affecting Wnt/beta-catenin signaling and supports chidamide plus azacitidine therapy in acute myeloid leukemia

**DOI:** 10.1101/2023.01.09.523194

**Authors:** Duanfeng Jiang, Qiuyu Mo, Haigang Shao, Jie Meng, Ruilan Zhong, Xunxiu Ji, Changjiu Liang, Wenyuan Lin, Fangping Chen, Min Dong

**Author notes:** **Correspondence author:** Min Dong, Department of Hematology, The second Affiliated Hospital of Hainan Medical University; Yehai Road, No. 368, Haikou, China.

## Abstract

**Background:** DACT gene is a potential Wnt antagonist and tumor suppressor gene. However, the expression of DACT gene in acute myeloid leukemia (AML) and its role are still unknown.

**Methods:** In this study, we first multidimensionally analyzed the expression of DACT gene in AML through RNA seq, qRT-PCR and Western blotting analysis as well as TCGA database analysis. Then DACT3 was identified as playing a critical role in AML, and its clinical significance was further explored. The molecular mechanism of DACT3 down-regulation in AML from aspects of DNA methylation and histone acetylation were inquired. Finally, the biological functions of DACT3 in AML were investigated by cell models *in vitro* and animal experiments *in vivo*.

**Results:** Compared with DACT1 and DACT2, DACT3 was the most differentially expressed DACT family member in AML, suggesting that DACT3 plays the major role. The mRNA and protein expression levels of DACT3 were significantly decreased and positively correlated in AML. Moreover, low expression of DACT3 is associated with no remission disease status and poor prognosis in AML patients. In addition, down-regulation of DACT3 in AML was not completely dependent on promoter methylation, but also affected by histone deacetylation. We found that loss of DACT3 in AML is related to activation of Wnt signaling pathway. Furthermore, DACT3 inhibits the growth of AML cells *in vitro* and in severe immunodeficiency (SCID) mice. Importantly, DACT3 improved the sensitivity of adriamycin in treating AML cells. Further research showed that co-treatment of chidamide and azacytidine up-regulates DACT3 expression and promotes cell apoptosis via inhibiting Wnt/β-catenin signaling in AML, suggesting a promising therapeutic prospects in FLT3-mutant AML.

**Conclusion:** This is the first study revealing the epigenetic regulatory molecular mechanism of the down regulated expression of DACT3 in AML. Our data also shows that DACT3 is a candidate gene for therapeutic target and targeting DACT3 has the potential to validate the combination therapy of DNMT inhibitors and HDAC inhibitors.

## Introduction

Acute myeloid leukemia (AML) is a hematopoietic stem/progenitor cell malignancy that remains the most common type of leukemia in adults [1]. Despite the application of intensive chemotherapy and hematopoietic stem cell transplantation (HSCT), with more than half of young patients and about 80% of old patients ultimately dying from AML [1,2]. AML is a heterogeneous disease caused by a variety of genetic alterations in epigenetic regulators, transcription factors, and signaling molecules [3]. A major group of AML-specific mutations encode epigenetic regulators including regulators of histone and DNA methylation such as MLL, EZH2, TET2, DNMT3A and IDH1/2 [4,5].

The classical Wnt/β-catenin signaling pathway is a major signal system of stem/progenitor cells and tumor cells [6]. Wnt binds to cell surface receptors, including frizzled (FZD) protein and LDL receptor related protein 6 (LRP6), and starts a transcription coactivator β-catenin is a stable transduction cascade, and then enters the nucleus to form a transcription complex with T cell factor (TCF) or lymphoid enhancer factor (LEF) to activate the expression of Wnt target gene c-Myc and cyclin D1 [7]. The aberrant Wnt/β-catenin signaling pathway exerts crucial roles in tumorigenesis, including regulating tumor stem cell renewal, cell proliferation, differentiation and invasion [8]. Increasing evidence demonstrates that dysregulation of the Wnt/β-catenin cascade contributed to the development and progression of hematological malignancies [9,10]. Activation of the Wnt/β-catenin pathway has been shown to be crucial to the establishment and maintenance of leukemic stem cells (LSCs), and its imbalance contributes to the relapse and development of AML [11,12]. It has been confirmed that Wnt signaling are abnormally activated and show a prognostic significance in AML [13,14]. β-catenin is highly expressed in most AML cells, and more higher in bone marrow LSCs compared to normal CD34^+^ cells. Moreover, the overexpression of β-catenin correlates with inferior patient survival in AML [15,16]. LEF-1 drives aberrant β-catenin nuclear localization in myeloid leukemia cells and its high expression in AML is associated with poor prognosis [17]. Moreover, many Wnt pathway components have a prognostic value in AML, including phosphorylated-GSK3β, sFRP2, PSMD2, et al [12,13 18,19]. Wnt/β-catenin signaling pathway plays an important role in the regulation of leukemic stem cells and leukemogenesis in AML [20], however, the molecular mechanism remains unclear.

Dishevelled-binding antagonist of beta-catenin (DACT), a β-catenin antagonist, plays a role by binding with the protein Dishevelled. With other names appeared successively in the literature, Dapper (Xenopus laevis), Frodo (zebrafish) and HDPR (human Dapper protein), it is now uniformly named DACT [21,22]. Three DACT gene family members are expressed during embryonic development and in the adult brains of mice [23]. The members of DACT protein family were initially identified by binding to Dishevelled (DVL, or Dsh), which is the core cytoplasmic protein of Wnt signaling [21,24]. DACT protein is a newly discovered Wnt signaling inhibitor, that it can bind to PDZ domain of DVL through C-terminal PDZ binding motif, promote the degradation of Dishevelled and antagonize Wnt signaling [22,24]. Recent studies have shown that DACT gene family members are often down regulated in a variety of solid tumors and are related to tumor progression. Mechanisms of DACT down-regulation are proved to be associated with epigenetic regulation in these tumors, including promoter methylation and histone modification. Furthermore, DACT is recognized as a tumor suppressor of Wnt/β-catenin pathway [25-27]. However, the role and epigenetic regulation mechanism of DACT family members are different in various tumors. For example, DACT1 is often down regulated related to methylation in gastric cancer, and DACT2 promoter is hypermethylated in lung cancer [28,29]. Unlike other Wnt signal antagonists silenced by DNA methylation, the epigenetic inhibition of DACT3 in colorectal cancer is mainly related to bivalent histone modification [30]. New evidence suggests that DACT1 and DKK1 expression were down regulated in AML patients, while they were highly expressed in ALL patients [31]. However, the role of DACT gene family members in AML remains unknown.

Epigenetic dysregulation has been proven to play a crucial role in progression of some malignancies, especially true of AML. The silencing of tumor suppressor genes in leukemia is mainly related to epigenetic modification, including methylation of CpG island in promoter region and bivalent histone modification [31,32]. Wnt signaling pathway is regulated by epigenetic regulation in AML, which is manifested as abnormal methylation of Wnt antagonist [33,34]. Over 64% of AML patients have abnormal methylation of Wnt antagonists, including sFRP1, sFRP2, DKK1 or DKK3, which is associated with poor prognosis [35]. Treatment with demethylated drug descitabine can restore expression of methylated Wnt antagonist and inactivation of Wnt pathway in AML [35,36]. Numbers of drugs used to treat AML has increased unprecedentedly in the past decade [1]. Most of them can be classified as targeted epigenetic therapy drugs, such as targeted inhibitors targeting LSD1, DOT1L, EZH2, BET/BRD4, IDH1/2, MEN1-MLL, etc. [37-39]. Decitabine (DAC) and azacytidine (AZA) are two major hypomethylating agents (HMAs), which have recently emerged as effective drugs for the treatment of AML. Even if success has been achieved in blood tumors, it is still controversial whether the epigenetic effects of HMAs can explain all or part of the therapeutic response [40]. In addition, histone deacetylase (HDAC) inhibitors do not seem to react as a single drug in AML. Therefore, the combination of HMAs and HDAC inhibitors has become a research hotspot in recent years [41,42]. Epigenetic therapies, including HMAs and HDAC inhibitors, were developed to reprogram the epigenetic abnormalities in AML. However, the molecular mechanisms and therapeutic effects of the two agents alone or their combination remain unknown.

Herein, we report that DACT3 is frequently downregulated in AML, and inhibits leukemia cell survival. Decreased DACT3 expression is partially caused by the DNA methylation. In addition, low DACT3 is associated with poor outcome in AML patients in our study. Moreover, we evaluated the effects of the hypomethylating agent azacytidine and HDAC inhibitors chidamide on apoptosis, and the potential of therapeutic strategies involving DACT3 targeting in AML cells. In particular, we aimed to elucidate the mechanisms underlying the effects of these epigenetic therapies on AML cell death. These findings further indicate that DACT3 might be a potential target for the treatment of AML.

### Patients and samples

A total of 142 AML patients (129 de novo AML and 13 relapsed/refractory AML) and 51 AML patients with complete remission (CR) were enrolled in this study approved by the Institutional Ethics Committee of the Third Xiangya Hospital of Central South University, from February 2017 to December 2019. The diagnosis and classification of patients are based on the French American British (FAB) classification [43] and the 2016 World Health Organization (WHO) criteria [44]. Relapse and CR are defined according to the recommendations of the European Leukemia Network (ELN) [45]. Control samples (n = 31) were from donors without any malignant BM disease, including 16 healthy donors of allogeneic transplantation, 12 iron deficient anemia (IDA) patients and 3 primary immune thrombocytopenia (ITP) patient [46]. BM mononuclear cells were enriched by Ficoll gradient purification, and the isolated mononuclear cells were directly frozen at -80 °C or cell lysate with Trizol was added. According to the Helsinki Declaration, informed consent was obtained, and the use of BM samples was approved by the Medical Ethics Committee of the Third Xiangya Hospital of Central South University.

### Cell lines and cell culture

Human myeloid leukemia cell lines K562, K562/ADR, THP1 and HL60 cells, human embryonic kidney 293 T cells, and human normal hematopoietic cell line GM12878 were purchased from the Cancer Institute of Central South University. The human lymphocytic leukemia cell line Jurkat was purchased from the Cell Bank of the Chinese Academy of Sciences (Shanghai). MLM13 and FLT3-ITD^+^ MV4-11 [47] cell lines were presented by Professor Zeng Hui from the First Affiliated Hospital of Jinan University. All leukemia cells were maintained in RPMI 1640 medium (Gibco, Australia), 293 T cells were cultured in RPMI DMEM medium (containing 10% fetal bovine serum, Gibco), and placed in a 5% CO_2_ humidified incubator at 37 °C. Adriamycin was added (final concentration of 1 μmol/L) to the complete culture medium of K562/ADR until 2 weeks before experiments [46].

### Chemical inhibitors treatment

The hypomethylating agent azacytidine (MedChemExpress) was dissolved in DMSO (0 to 10 μM in culture). HDAC inhibitors chidamide (Selleck) was dissolved in DMSO (0.5 to 5 μM in culture in culture). The anthracycline agent adriamycin (Sigma-Aldrich) was dissolved in DMSO (1 μM in culture). Azacytidine and chidamide were used to treat HL60 and THP1 cells for indicated and cells were harvested for cell apoptosis analysis, qRT-PCR and western blotting.

### RNA-seq assay

Total RNA of cells was extracted according to manual of TRIzol reagent (Invitrogen, USA). After analyzing RNA sample quality, BGI technology (Beijing, China) was used to synthesize and sequence the cDNA libraries [48,49]. Library preparation and transcriptome sequencing was performed on an Illumina platform by Annoroad Gene Technology (Beijing, China) [50,51]. Differentially expressed genes (DEGs) were identified using the DESeq2 R package. The criteria for DEGs was set up as fold change (FC, log2) >2 or <−2, Q-value <0.05, and FDR<0.05 [46,48].

### DNA isolation, bisulfite modification and RQ-MSP

Genomic DNA was extracted from cells using Genomic DNA Isolation Kit (Vazyme, China; #DC112), then modified using the EpiArt™ DNA Methylation Bisulfite Kit (Vazyme, #EM101). Genomic DNA isolation and modification were performed as reported previously. Real-time quantitative methylation-specific PCR (RQ-MSP) was applied to examine DCT3 methylation level using ChamQ Universal SYBR qPCR Master Mix (Vazyme, #Q711) with primers presented in Supplementary Table S1. RQ-MSP conditions for DACT3 and other genes methylation level detection were 95 °C for 5 min, 40 cycles for 10 s at 95 °C, 30s at 63 °C (collect fluorescence), and finally followed by the melting program at 95 °C for 15 s, 60 °C for 60 s, and 95 °C for 15 s. The gene ALU was used as the endogenous control to calculate DACT3 methylation level [52]. Normalized DACT3 methylation level (N_M-DACT3_) was calculated by 2^−ΔΔCT^ methods [53]. N_M_-DACT3 was calculated using the following formula: N_M-DACT3_ = (E_M-DACT3_)^ΔCT M-DACT3 (control-sample)^/(E_ALU_) ^ΔCT ALU(control-sample)^.

### Plasmid and virus production

The full-length cDNA of human DACT3 was synthesized by HonorGene (Changsha, China) and cloned into the lentiviral expression vector pHG-CMV. Two small hairpin RNA (shRNA1 and shRNA2) targeting human DACT3 were provided by Tsingke Biotechnology (Beijing, China) and were cloned into pLKO-tet-on lentiviral vector. For stable transfection assays, all vectors were packaged into lentiviruses with packaging plasmids psPAX2 and pMD2G in HEK-293 T cells [54]. The lentiviruses were transfected into HL-60 and THP1 cells, and stably transduced overexpression (OE) and knockdown cells were selected with 2 μg/ml puromycin (Sigma, USA). All transfection experiments were performed using the INVI DNA/RNA Transfection Reagent (Invigentech, USA) according to the manufacturer’s instructions. After 48 hours of transfection, the cells were collected for qRT-PCR or western blotting analysis to confirm transfection efficacy [55]. The primer sequences were listed in Supplementary Table S2.

### Delivery of siRNA and cell transfection

The siRNA targeting β-catenin and control siRNA were purchased from Tsingke Biotechnology (Beijing, China). The HL-60 and THP1 cells were transfected with siRNA using the INVI DNA/RNA Transfection Reagent (Invigentech, USA) according to the manufacturer’s instructions. The transfection efficiency was assessed by qRT-PCR. The sequences of siRNA were listed in Supplementary Table S2.

### Dual-luciferase reporter assay

The stably transduced DACT3 overexpression (OE) or knockdown cells were seeded into 96-well plates and cotransfected with the TOP/FOP flash reporter plasmid, pRL-TK reporter plasmid, and β-catenin siRNA or negative control siRNA using INVI DNA/RNA Transfection Reagent (Invigentech, USA). After transfection for 48 hours, cells were harvested for relative luciferase activity analysis using the Dual-Luciferase Reporter Assay System (US Everbright, USA) following the manufacturer’s instructions. Luciferase activity was measured with a luminometer (Lumat LB9507, Berthold, Germany). Results of triplicate transfections were normalized to Renilla luciferase activity.

### Immunofluorescence analysis

Immunofluorescence analysis was performed according to standard protocols as described previously [56]. Cells were collected and placed on glass substrates and fixed with 4 % paraformaldehyde for 20 minutes. After permeabilized and blocked, the fixed cells were incubated with primary antibody against β-catenin (Sigma-Aldrich, USA) and secondary antibody conjugated with Alexa Fluor 488 goat anti-rabbit (Affinity Biosciences, USA), followed by counterstaining with 4′,6-diamidino-2-phenylindole (DAPI) (Sigma, USA). The images were analysed and captured by confocal fluorescence microscope (Olympus, Japan).

### The cancer genome atlas (TCGA) gene expression data analysis

The gene expression differences between AML patients and normal controls were conducted by Gene Expression Profiling Interactive Analysis (GEPIA) network platform (http://gepia.cancer-pku.cn/detail.php) [46,57]. Gene expression data of AML patients were derived from The Cancer Genome Atlas (TCGA, http://www.cancergenome.nih.gov). The mRNA expression of target genes were compared between AML cases (n = 173) and the normal controls (n = 70).

### Survival analysis

Gene expression levels and clinical survival information of 162 AML patients, were retrieved from Genomoscape dataset (http://www.genomicscape.com/microarray/survival.php.). All patients were stratified by DACT3 expression levels into different ranks, to categorize patients into either a high cohort or low cohort. Kaplan-Meier data of AML patients were used to analyze the overall survival (OS).

### In vivo model of acute myeloid leukemia

Animal studies were approved by the Institutional Animal Care and Use Committee of Central South University, China. Female B-NDG mice (18–20g, 5-7 weeks old; Jiansu Biocytogen Co., Ltd., Nantong, China) were used [56,58]. AML xenograft mouse model was established through subcutaneous injection of 5 × 10^6^ HL60 cells (n = 20 mice), DACT3 shRNA HL60 cells (sh-DACT3, n = 5 mice) and HL60 cells transfected with control shRNA (NC, n = 5 mice). After 10 days, when the tumours became palpable, the 20 mice that injection of HL60 cells were randomised into four groups: vehicle, chidamide, azacytidine, and combination groups. Tumour-bearing mice were treated with chidamide (10 mg/kg) [59] and/or azacytidine (0.5 mg/kg) injected intraperitoneal daily for the first 5 days of every 14 day treatment cycle [60], PBS injected for the vehicle group. Treatment was continued for the entire duration of the study and mice were killed before tumors volume exceeded 2000 mm^3^. Tumor volume determined by measurements obtained from digital caliper and calculated as V = 0.5 × length × width^2^. After mice were sacrificed, tumors were excised and weighed. Survival was also followed and presented with a Kaplan-Meier survival plot.

### Statistical analyses

Data are presented as mean ± standard deviation (SD). Statistical analyses were conducted using SPSS software version 19.0 and GraphPad Prism 7.0. Unpaired Students’ t-test or Mann Whitney U test were carried to compare the difference of continuous variables between two groups, whereas Pearson chi-square analysis/Fisher exact test was applied to compare the difference of categorical variables between two groups. Correlation analysis was performed by Spearman test. The χ^2^ analysis was applied to compare the proportions of the two groups of variables. The paired Wilcoxon was conducted to analyze the difference before and after treatment. Survival was presented with a Kaplan-Meier survival plot and calculated using log-rank tests [61]. P value < 0.05 was considered to indicate statistical significance.

A detailed description of materials and methods not shown is available in the *Supplementary Information*.

## Results

### DACT3 expression is repressed in primary acute myeloid leukemia samples and cell lines

The DACT gene families (DACTs) has three members, including DACT1, DACT2, and DACT3 [62]. To investigate the role of DACTs in AML, we first detected the transcription level of the genes using RNA-seq analysis. Hierarchical clustering analysis showed that DACT3, a Wnt signaling inhibitor [30], was significantly down regulated in AML patients compared with normal controls, whereas DACT1 and DACT2 were not of the downregulated genes (Figure 1A). These data showed that compared with DACT1 and DACT2, DACT3 may play the major role in AML. Consistently, DACTs mRNA expression (RNA-Seq) data from TCGA showed that DACT3 expression was decreased (P<0.01) in a cohort of 173 AML patients compared with 70 normal controls, whereas DACT1 and DACT2 expression have no differences (P>0.05) between AML patients and normal controls (Figure 1B). More importantly, qRT-PCR analysis showed that DACT3 mRNA expression was reduced in 142 AML samples (P<0.001), whereas DACT1 and DACT2 expression levels did not show significant differences between tumor and normal controls (P=0.53 and P=0.22, respectively) (Figure 1C). These results indicated that DACT3, not other DACT family gene, plays a crucial role in AML.

**Figure 1.**
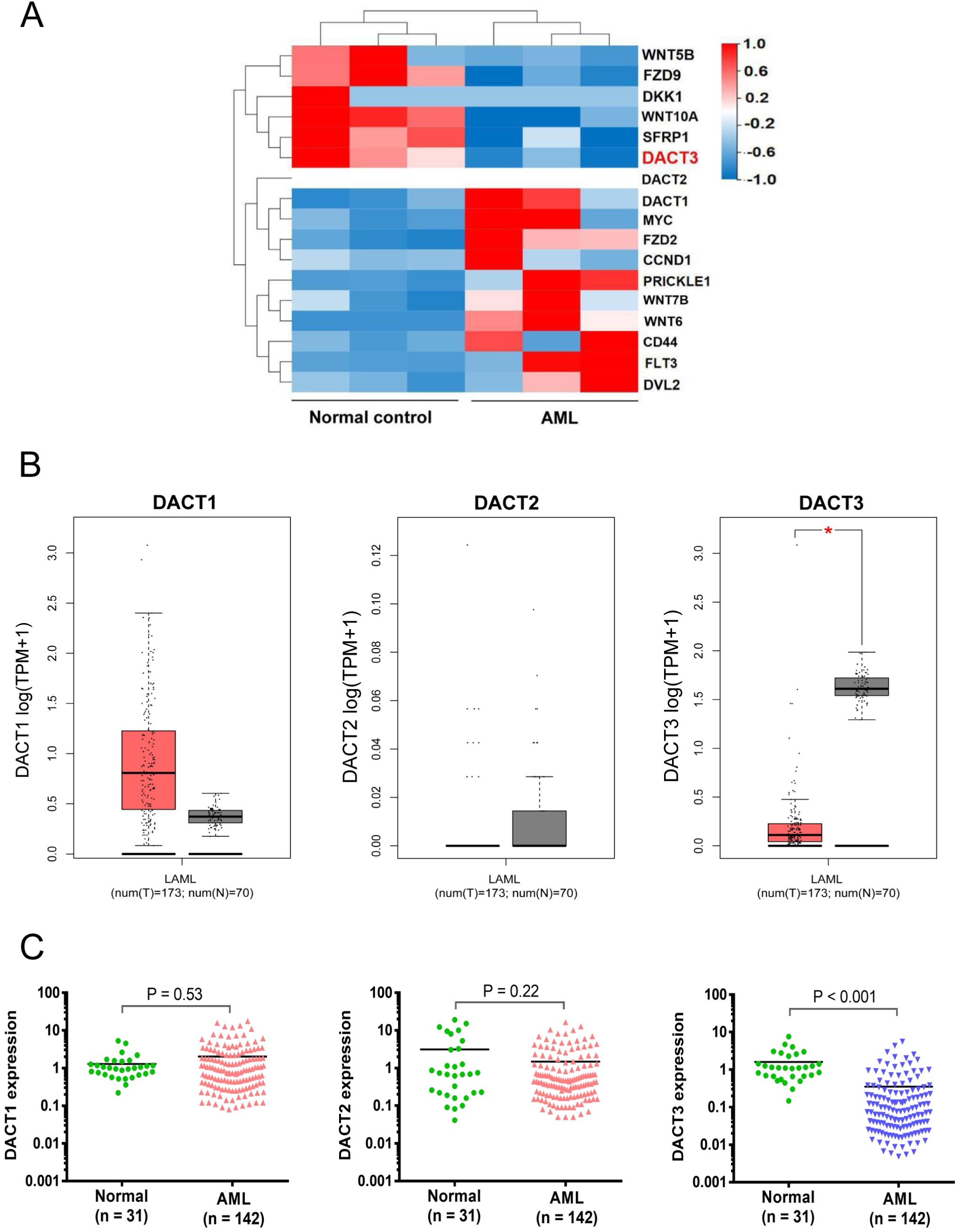
Loss of DACT3 mRNA expression in AML patients. (**A**) RNA-seq analysis of DACT3 expression in AML patients (n=4) and normal controls (n=3). Hierarchical cluster analysis of the differentially expressed genes (DEGs) including 12 Wnt signaling components, CD44, FLT3 and DACT gene families (DACTs). Upregulated genes are shown in red and downregulated genes are shown in blue. (**B**) RNA-seq mRNA expression data from TCGA database were used to compare DACT3 expression between AML patients (n = 173) and normal contrals (n = 70). ^*^P<0.01. (**C**) qRT-PCR analysis of DACTs mRNA levels in primary AML samples (n = 142) and normal contrals (n = 31).

The above results confirmed that the mRNA level of DACT3 was significantly reduced in AML. To explore the protein level of DACT3 in AML, western blotting was used to detect the expression of DACT3 in bone marrow mononuclear cells of AML patients. As shown in Figure 2A-D, DACT3 protein level was significantly reduced (P<0.001) in 18 AML patients compared with 6 normal controls. In addition, both mRNA (Figure 2E) and protein (Figure 2F and G) of DACT3 were significantly lower expressed (P<0.05) in AML cell lines THP1, K562, K562/ADR, HL60, MOLM13 and MV4-11, compared with normal hematopoietic cell GM12878. Furthermore, correlation analysis (R^2^=0.754, P=0.024) indicated that the mRNA expression level and protein level of DACT3 were positively correlated in AML cell lines (Figure 2H). Interestingly, the mRNA expression of DACT3 in adriamycin resistant cell line K562/ADR was significantly lower than that in the parent cell line K562. To sum up, the expression of DACT3 in AML cells is generally reduced, and the loss of DACT3 may be associated with adriamycin resistant. These data suggested that DACT3 may be a potential molecular target for diagnosis and treatment of AML.

**Figure 2.**
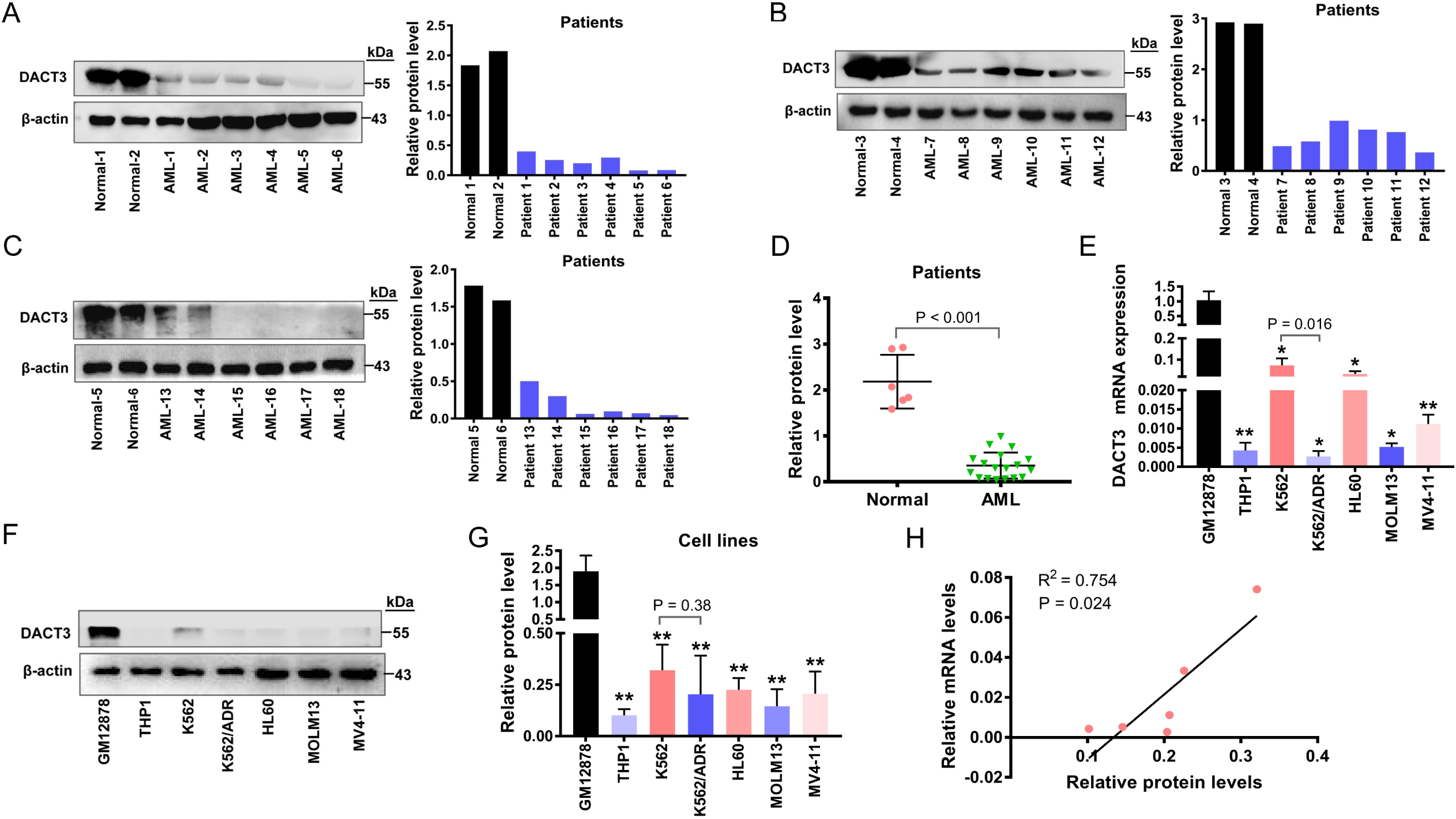
Downregulation of DACT3 protein level in AML patients and cell lines. (**A, B** and **C**) The expression of DACT3 protein was detected in bone marrow mononuclear cells of 6 normal controls and 18 AML patients. (**D**) statistical analysis of the DACT3 protein levels between AML and normal controls. (**E**) qRT-PCR analysis of DACT3 mRNA expression from the indicated AML cell lines. (**F** and **G**) Western blotting analysis of DACT3 in AML cell lines. (**H**) Pearson correlation analysis for DACT3 mRNA and protein expression in the 6 AML cell lines (Pearson r = 0.754, P = 0.024).

### Low DACT3 expression is correlated with disease status, age, white blood cells (WBCs), BM blasts, FLT3 and DNMT3A mutations in acute myeloid leukemia patients

It is worth noting that, lower DACT3 expression was observed in patients with newly diagnosed AML (n = 129, P < 0.001) and relapsed AML (n = 13, P < 0.001) than in patients with complete remission (n = 51, Figure 3A). In addition, DACT3 mRNA levels were determined in 10 patients at the time of being newly diagnosed, complete remission and relapse. A common feature was that the expression of DACT3 was low at new diagnosis, increased after complete remission (P = 0.026, Figure 3B), and increased again at relapse (P = 0.025, Figure 3B). To explore the correlation of DACT3 expression with clinical features in AML patients, we divided the patients into a low DACT3 expression group (DACT3^low^, the first half, n=64) and a high DACT3 expression group (DACT3^high^, the second half, n=65) according to the cut-off value of 0.064 (median DACT3 expression level). The comparisons of clinical features and laboratory parameters between the two groups were shown in Table 1 and Table 2. The low expression of DACT3 was found to be associated with higher white blood cells (WBCs) (P=0.036), higher BM blasts (P=0.019), FLT3 mutation (P=0.040) and DNMT3A mutation (P=0.043) (Table 1). However, we did not observe significant differences in sex, hemoglobin (HB), platelets (PLT), FAB classifications, karyotypes, ELN risk stratification and cytogenetic risk between DACT3^low^ and DACT3^high^ patients (Table 1).

**Table 1.** Correlation of DACT3 expression with clinical parameters in AML patients.

| Patient's parameters | Total<br>(n=129) | Status of DACT3 expression |  | P value |
| --- | --- | --- | --- | --- |
|  |  | Low (n=64) | High (n=65) |  |
| Sex, male/female | 77/52 | 40/24 | 37/28 | 0.518 |
| Median age, years (range) | 51 (12-81) | 48 (13-77) | 54 (12-81) | 0.058 |
| Median WBC,<br>×10<sup>9</sup>/L (range) | 15.6 (0.3-381.6) | 21.3 (0.8-381.6) | 12.4 (0.3-210.9) | 0.036 |
| Median hemoglobin,<br>g/L (range) | 73 (33-151) | 70 (33-132) | 75 (39-151) | 0.658 |
| Median platelets,<br>×10<sup>9</sup>/L (range) | 30 (2-984) | 27 (2-400) | 33 (4-984) | 0.345 |
| Median BM blasts %, (range) | 73.0 (16.0-97.0) | 75.0 (24.0-97.0) | 69.5 (16.0-95.5) | 0.019 |
| FAB subtypes (%) |  |  |  | 0.261 |
| M0 | 1 (0.8) | 0 (0.0) | 1 (1.5) |  |
| M1 | 13 (10.1) | 8 (12.5) | 5 (7.7) |  |
| M2 | 49 (38.0) | 30 (46.8) | 19 (29.2) |  |
| M3 | 17 (13.2) | 7 (10.9) | 10 (15.4) |  |
| M4 | 13 (10.1) | 6 (9.4) | 7 (10.8) |  |
| M5 | 34 (26.4) | 12 (18.8) | 22 (33.9) |  |
| Not determined | 2 (1.6) | 1 (1.6) | 1 (1.5) |  |
| ELN risk stratification |  |  |  | 0.158 |
| Favorable | 50 (38.8) | 20 (31.3) | 30 (46.2) |  |
| Intermediate | 44 (34.1) | 24 (37.5) | 20 (30.8) |  |
| Poor | 19 (14.7) | 13 (20.3) | 6 (9.2) |  |
| No data | 16 (12.4) | 7 (10.9) | 9 (13.8) |  |
| Cytogenetic risk (%) |  |  |  | 0.227 |
| Favorable | 39 (30.2) | 16 (25.0) | 23 (35.4) |  |
| Intermediate | 59 (45.7) | 32 (50.0) | 27 (41.5) |  |
| Poor | 15 (11.6) | 10 (15.6) | 5 (7.7) |  |
| No data | 16 (12.4) | 6 (9.4) | 10 (15.4) |  |
| Karyotypes (%) |  |  |  | 0.248 |
| t(8;21)/RUNX1-RUNX1T1 | 18 (14.0) | 13 (20.3) | 5 (7.7) |  |
| inv(16)/CBFβ-MYH11 | 4 (3.1) | 2 (3.1) | 2 (3.1) |  |
| t(15;17)/PML-RARA | 17 (13.2) | 7 (10.9) | 10 (15.4) |  |
| 11q23/MLL | 7 (5.4) | 3 (4.7) | 4 (6.1) |  |
| Normal karyotype | 41 (31.8) | 24 (37.5) | 17 (26.2) |  |
| Complex karyotype | 5 (3.9) | 2 (3.1) | 3 (4.6) |  |
| Other karyotype | 21 (16.3) | 8 (12.5) | 13 (20.0) |  |
| No data | 16 (12.4) | 5 (7.8) | 11 (16.9) |  |
| FLT3 (%) |  |  |  | 0.040 |
| FLT3-ITD | 16 (12.4) | 12 (18.8) | 4 (6.2) |  |
| FLT3-TKD | 4 (3.1) | 0 (0.0) | 4 (6.2) |  |
| wild | 85 (65.9) | 40 (65.6) | 45 (66.1) |  |
| No data | 24 (18.6) | 12 (15.6) | 12 (21.5) |  |
| CEBPA |  |  |  | 0.210 |
| Single Mutation | 3 (2.3) | 2 (3.1) | 1 (1.5) |  |
| Double Mutation | 15 (11.6) | 11 (17.2) | 4 (6.2) |  |
| wild | 87 (67.4) | 39 (60.9) | 48 (73.8) |  |
| No data | 24 (18.6) | 12 (18.8) | 12 (18.5) |  |
| DNMT3A |  |  |  | 0.043 |
| mutated | 6 (4.7) | 0 (0.0) | 6 (9.2) |  |
| wild | 99 (76.7) | 52 (81.2) | 47 (72.3) |  |
| No data | 24 (18.6) | 12 (18.8) | 12 (18.5) |  |
| IDH1 |  |  |  | 0.934 |
| mutated | 7 (5.4) | 3 (4.7) | 4 (6.1) |  |
| wild | 98 (76.0) | 49 (76.5) | 49 (75.4) |  |
| No data | 24 (18.6) | 12 (18.8) | 12 (18.5) |  |
| IDH2 |  |  |  | 0.402 |
| mutated | 10 (7.8) | 4 (6.2) | 6 (9.2) |  |
| wild | 95 (73.6) | 48 (75.0) | 47 (72.3) |  |
| No data | 24 (18.6) | 12 (18.8) | 12 (18.5) |  |
| NPM1 (%) |  |  |  | 0.982 |
| mutated | 23 (17.8) | 11 (17.2) | 12 (18.5) |  |
| wild | 82 (63.6) | 41 (64.0) | 41 (64.0) |  |
| No data | 24 (18.6) | 12 (18.8) | 12 (18.5) |  |

**Table 2.** Patients’ information.

| Patient's characteristics | Normal control<br>(n=31) | AML-CR<br>(n = 51) | AML-Relapse<br>(n = 13) |
| --- | --- | --- | --- |
| Median age in years (range) | 32 (19-50) | 43 (18-81) | 51 (20-66) |
| Sex (male/female) | 18 (56.2%)/14 (43.8%) | 28 (54.9%)/23 (45.1%) | 4 (30.8%)/9 (69.2%) |
| Unfavorable fusion gene | - | 3 (8.6%) | 5 (38.5%) |
| Unfavorable karyotype | - | 3 (8.6%) | 4 (30.8%) |

**Figure 3.**
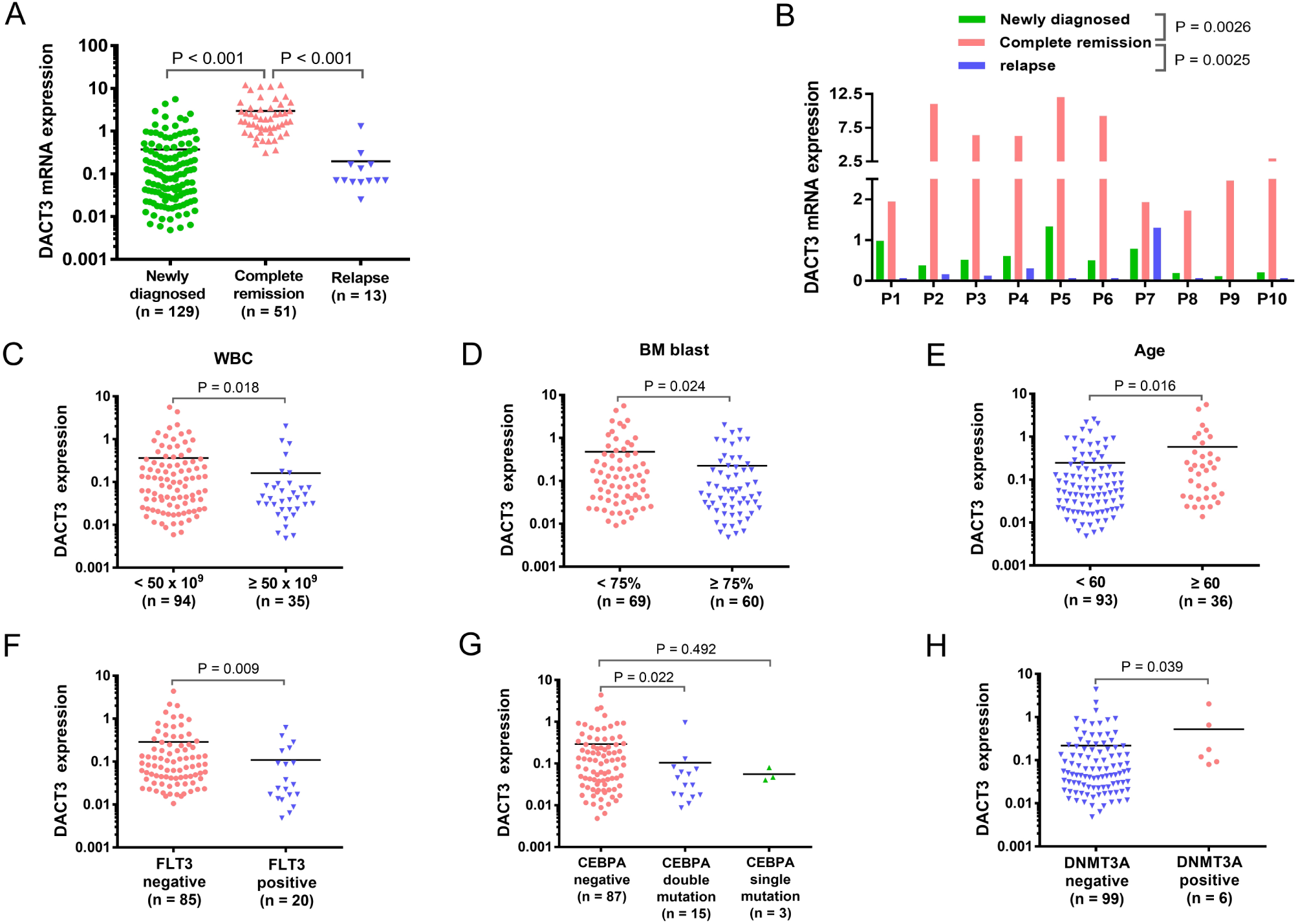
Correlation analysis between DACT3 expression and clinical parameters in AML patients. (**A**) mRNA expression level of DACT3 in bone marrow samples of AML patients with newly diagnosis (n=129), complete remission (n=51) and relapse (n=13) using qRT-PCR. (**B**) qRT-PCR analysis of DACT3 mRNA expression in 10 AML patients at their stage of newly diagnosis, complete remission and relapse. DACT3 expression difference between patients with peripheral WBC count less and more than 50 × 109/L (**C**), bone marrow blasts less and more than 75% (D), age less and more than 60 years (**E**), with and without gene mutations [FLT3 (**F**), CEBPA (**G**) and DNMT3A (**H**)]. WBC: white blood cells; BM: bone marrow.

Interestingly, we found that the expression of DACT3 was lower in AML patients with WBCs ≥50 × 10^9^/L than in those with WBCs <50 × 10^9^/L (P=0.018, Figure 3C), and lower in AML patients with BM blasts ≥75% than in those with BM blasts <75% (P=0.024, Figure 3D). While the expression of DACT3 was higher in AML patients older than 60 years than in AML patients younger than 60 years (P=0.016, Figure 3E). Consistent with the previous results, the low expression of DACT3 was associated with FLT3 mutation (P=0.009, Figure 3F) and CEBPA double mutation (P=0.022, Figure 3G), other than with CEBPA single mutation (P=0.492, Figure 3G). It is worth noting that patients with DNMT3A mutations show high expression of DACT3 (P=0.039, Figure 3H), which may indirectly suggest that the expression of DACT3 is affected by DNA methylation. Because gene mutation reduces the activity of DNMT3A which can induce CpG hypomethylation and affect the expression of genes [63].

### Low DACT3 expression is associated with poor survival in patients with acute myeloid leukemia

In this study, 129 patients who can be evaluated were received median follow-up period of 11 months (1-27 months). Kaplan-Meier survival analysis showed that DACT3^low^ patients had significantly shorter overall survival (OS) (P=0.034, Figure 4A) than DACT3^high^ patients in the whole-cohort AML. Among the 112 non-M3 AML group, DACT3^low^ cases (n=55) also showed significantly shorter OS and event-free survival (EFS) than DACT3^high^ cases (n=57) (OS: P=0.034, Figure 4C; EFS: P = 0.036, Figure 4D). Although there were no significant difference in EFS between DACT3^low^ and DACT3^high^ patients of whole-cohort AML (P=0.265, Figure 4B) and cytogenetically normal AML (CN-AML) (P=0.198, Figure 4F), and no difference in OS between DACT3^low^ and DACT3^high^ patients of CN-AML (P=0.208, Figure 4E), a trend of separation may emerge as the extension of follow-up time. Furthermore, rank statistic AML patients from Genomoscape dataset at half, one-third and two-thirds quantile, showed all the OS of DACT3^low^ cases was shorter than those of DACT3^high^ cohort (P=0.0076, 0.013 and 0.028, respectively; Figure 4G-I).

**Figure 4.**
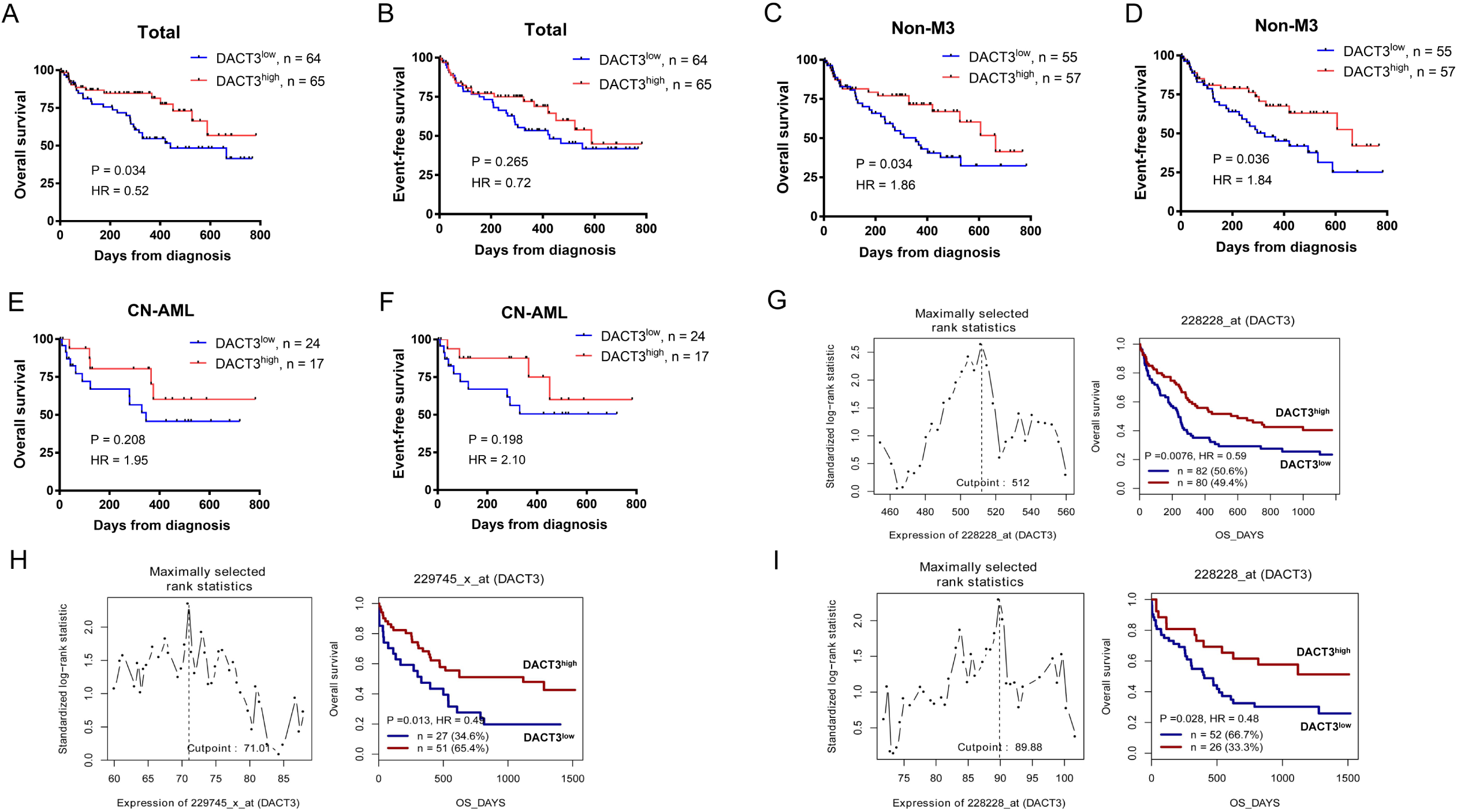
Low expression level of DACT3 is related to poor survival of AML patients. (**A**-**F**) Survival analysis of AML patients in our study according to DACT3 expression: (**A**) Overall survival (OS) of whole cohort AML; (B) Event free survival (EFS) of whole cohort AML; (**C**) OS of non M3 AML; (**D**) EFS of non M3 AML; (**E**) OS of cytogenetically normal AML (CN-AML); (**F**) EFS of CN-AML. (**G**-**I**) Survival analysis of AML cases from GenomicScape data online according to DACT3 expression. In different probe groups, rank statistic AML patients at half (**G**), one-third (**H**) and two-thirds quantile (**I**), the low expression of DACT3 was related to wose OS of AML patients (P=0.0076, 0.013 and 0.028, respectively).

We then performed univariate analyses and multivariate analyses on OS in the total 129 AML patients, including the expression level of DACT3, age, WBC, cytogenetic risk, and NPM1, FLT3-ITD, CEBPA, DNMT3A, IDH1/IDH2 mutations (mutant vs. wild-type). As shown in Table 3, DACT3 expression was significantly and independently correlated with a worse OS both in univariate (P=0.038) and multivariate analysis (P=0.028). Besides, WBCs and cytogenetic risk were related to poorer OS both in univariate analysis (P=0.036; P=0.022; respectively).

**Table 3.** Results of univariate and multivariate analysis for overall survival in AML patients.

|  | Univariate |  | Multivariate |  |
| --- | --- | --- | --- | --- |
|  | HR (95% CI) | P value | HR (95% CI) | P value |
| DACT3 (high vs. low) | 0.507 (0.266-0.964) | 0.038 | 0.339<br>(0.129-0.891) | 0.028 |
| Age (>mid vs. ≤mid) | 1.817(0.985-3.354) | 0.056 | 1.796<br>(0.811-3.975) | 0.149 |
| WBC (>mid vs. ≤mid) | 1.958 (1.044-3.674) | 0.036 | 1.668<br>(0.696-3.997) | 0.252 |
| Cytogenetic risk (poor vs.<br>intermediate vs. favorable) | 1.751 (1.084-2.828) | 0.022 | 1.803<br>(0.987-3.294) | 0.055 |
| NPM1 (mutated vs. wild) | 1.419 (0.662-3.041) | 0.368 | 0.478<br>(0.114-1.995) | 0.311 |
| FLT3-ITD (pos vs. neg) | 1.965 (0.853-4.525) | 0.113 | 1.066<br>(0.263-4.320) | 0.928 |
| CEBPA (mutated vs. wild) | 0.806 (0.334-1.950) | 0.633 | 0.473<br>(0.173-1.288) | 0.143 |
| IDH1 (mutated vs. wild) | 2.396 (0.827-6.943) | 0.107 | 1.819<br>(0.479-6.908) | 0.380 |
| IDH2 (mutated vs. wild) | 1.667 (0.640-4.343) | 0.296 | 2.740<br>(0.728-10.321) | 0.136 |

### Down-regulation of DACT3 in acute myeloid leukemia is related to DNA methylation

Previously studies showed that DACT1 and DACT2 were regulated by DNA methylation in breast, esophageal and nasopharyngeal cancer [25,64,65]. DACT3 is histone modified in colorectal cancer as well as affected by demethylation drugs [30]. However, the regulatory mechanism of DACT methylation and histone modification in AML has not been reported. It’s well known that DNMT1, DNMT3A and DNMT3B are the main methyltransferases regulating DNA methylation; MeCP2 and MBD2 are the main methylation binding proteins regulating DNA methylation [66-69]. Therefore, we first investigated mRNA expression of these genes from TCGA and found that DNMT3A was significant higher in AML than in normal control (P<0.05, Figure 5A). Except that DNMT3B decreased in AML (P<0.05, Figure 5A), other methylation regulating genes showed an increasing trend in AML(Figure 5A and B). Then we confirm the mRNA expression of these genes in 129 patients with primary AML and 31 normal controls with qRT-PCR analysis. As shown in Figure 5C and D, the mRNA expression of DNMT1 (P=0.015), DNMT3A (P=0.047) and DNMT3B (P=0.030) in AML patients were higher than those in normal controls. This was consistent with the literature reports [67,70], suggesting that DNMTs may participate in DACT3 hypermethylation of AML. In addition, the previous results of this study (Figure 3H) showed that patients with DNMT3A mutation (if functional inactivation) showed higher expression of DACT3, which further confirmed that the expression of DACT3 was affected by DNMTs. The mRNA expression of MBD2 was significantly increased (P<0.001, Figure 5D), suggesting that methylation binding protein MBD2 might participate in the transcriptional inhibition of DACT3 in AML.

**Figure 5.**
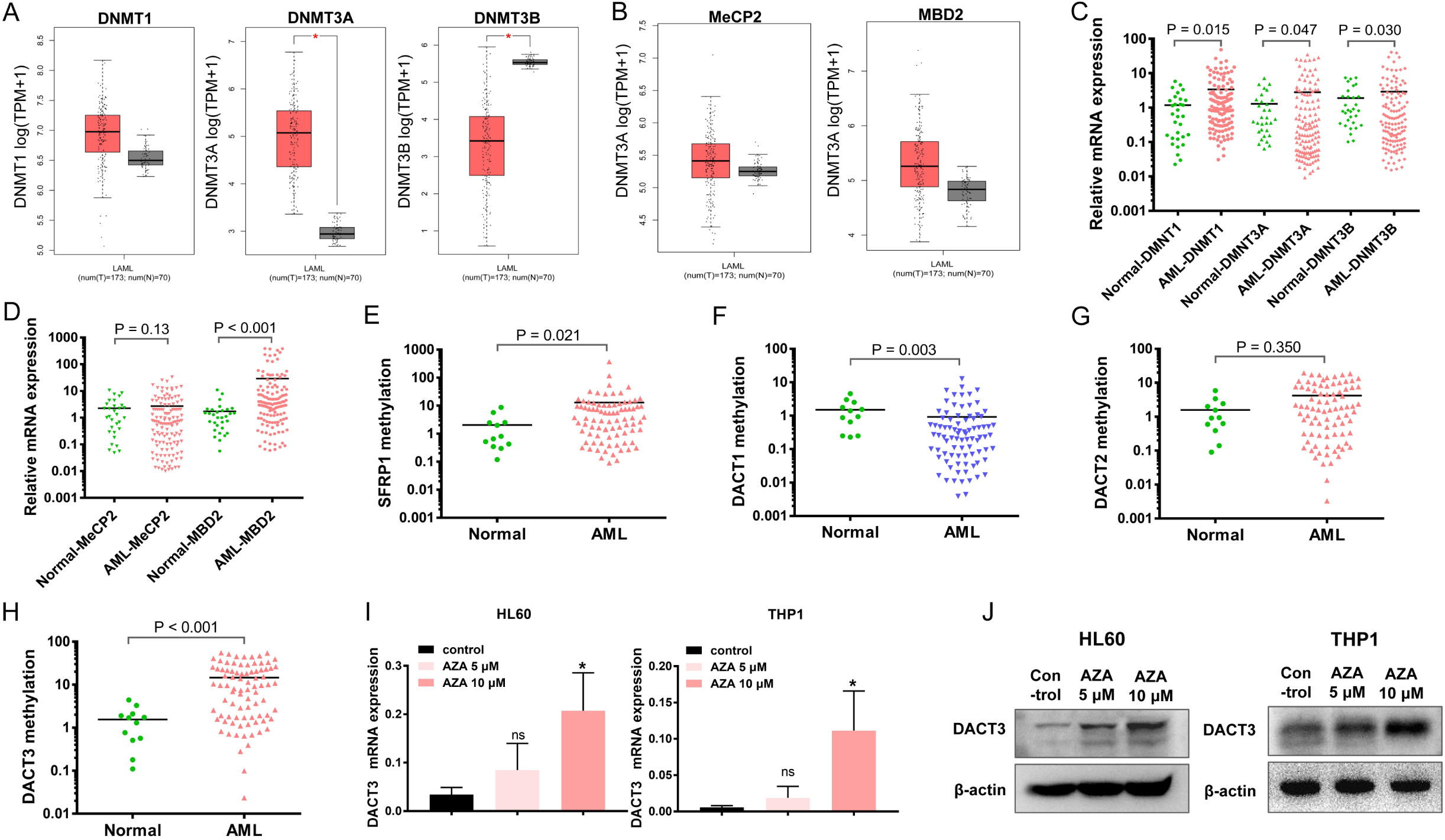
Loss of DACT3 in AML is associated with DNA methylation. RNA-seq mRNA expression data from TCGA database were used to compare (**A**) DNA methyltransferase (DNMT1, DNMT3A and DNMT3B) and (**B**) methylation binding protein (MeCP2 and MBD2) expression between AML patients (n = 173) and normal contrals (n = 70); ^*^P<0.05. (**C** and **D**) Analysis of DNMT1, DNMT3A, DNMT3B, MeCP2 and MBD2 mRNA levels between 129 newly diagnosed AML patients and 31 normal controls of our center. (**E**-**H**) Real-time quantitative methylation-specific PCR (RQ-MSP) analysis of SFRP1, DACT1, DACT2 and DACT3 methylation levels in 85 newly diagnosed AML patients and 12 normal controls. After treatment with different dose of azacytidine (AZA) (0, 5 and 10 μM) for 48 hours, the mRNA (**I**) and protein (**J**) expression levels of DACT3 were determined in AML cell lines HL60 and THP1; compared with control: ^ns^P>0.05; ^*^P <0.05.

Previous studies confirmed that the Wnt inhibitor SFRP1 is usually methylated in AML cells and was designed as a positive control for methylation analysis in this study. The methylation levels of SFRP1, DACT1, DACT2 and DACT3 in BM samples from 85 de novo AML patients and 12 normal controls were evaluated by real-time quantitative methylation specific PCR (RQ-MSP) [71]. The results showed that the median methylation level of the positive control gene SFRP1 (Figure 5E, P=0.021) and DACT3 (Figure 5H, P<0.001) were significantly higher in AML patients than that of normal controls, whereas the methylation level of DACT1 (Figure 5F; lower, P=0.003) and DACT2 (Figure 5G, P=0.352) were not higher than the normal controls. To investigate the effect of DNA methylation on DACT3 expression, AML cell lines HL60 and THP1 were treated with different concentrations of DNA methyltransferase inhibitor azacytidine for 48 hours. As shown in Figure 5I, azacytidine treatment upregulated the mRNA level of DACT3 in a dose-dependent manner. Consistently, the protein levels of DACT3 upregulated by azacytidine were also observed in western blotting analysis (Figure 5J). These results suggest that DACT3 is regulated by methylation in AML cells, which is consistent with the previously studies that DACT gene is regulated by promoter methylation in solid tumors [26,72,73].

### Expression of DACT3 in acute myeloid leukemia is affected by histone acetylation

To explore the effect of histone deacetylation on the expression of DACT3 in AML cells, HL60 and THP1 cells were treated with different dose of histone deacetylase inhibitor chidamide for 24 hours. As shown in Figure 6A and B, the mRNA expression of DACT3 increased after chidamide treatment, indicating that DACT3 expression in AML is affected by histone deacetylation. In order to further confirm this hypothesis, the AML cell lines HL60 and THP1 were treated with chidamide (for 24 hours) and azacytidine (for 48 hours) singly or in combination. As shown in Figure 6C and D, the combination treatment significantly increased the DACT3 mRNA expression in AML cells compared with the monotherapies (P<0.05). The results suggested that DACT3 expression in AML may also be affected by histone deacetylation.

**Figure 6.**
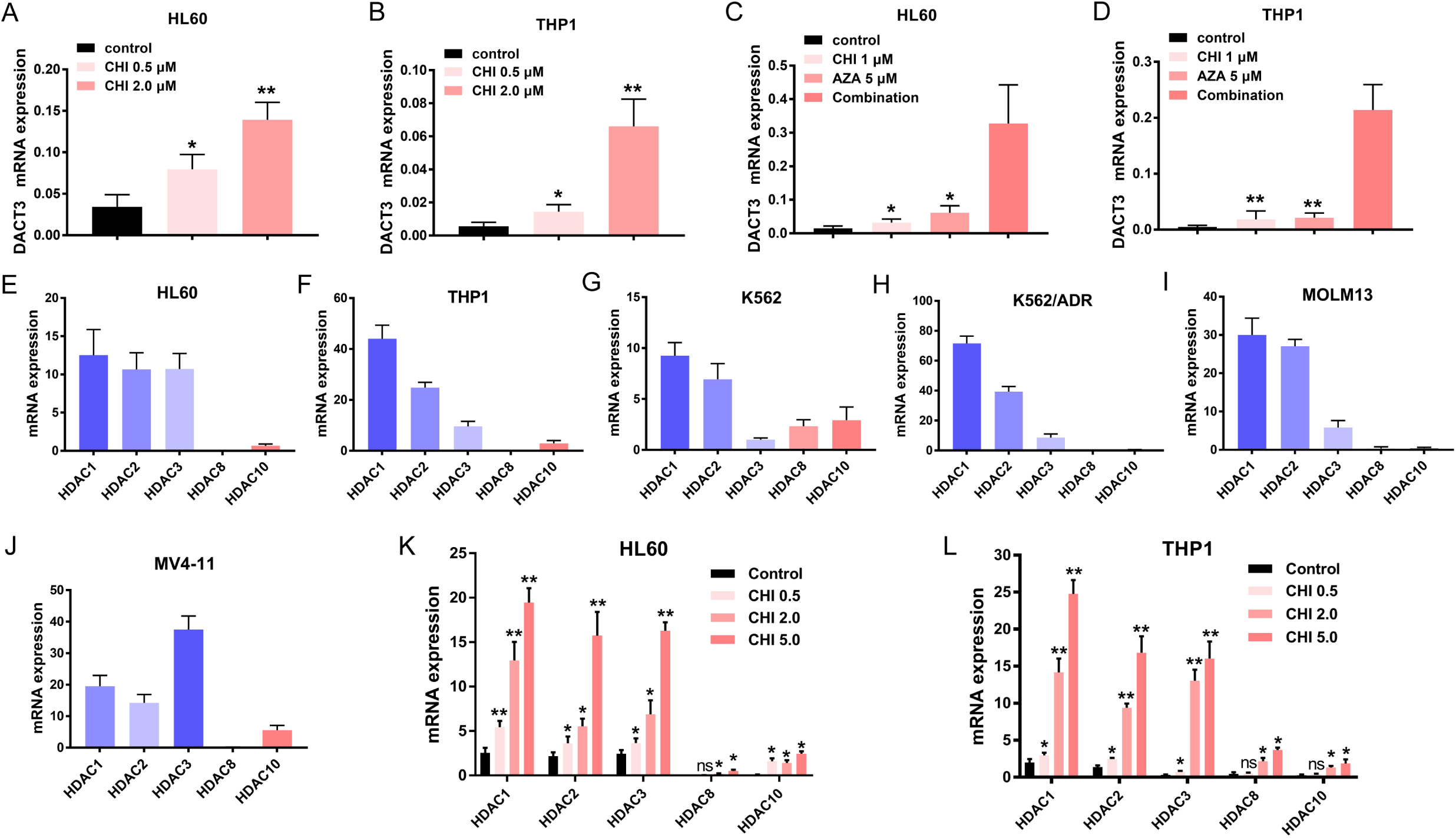
Expression of DACT3 in AML is affected by histone acetylation. After treatment with different dose of chidamide (CHI) (0, 0.5 and 2.0 μM) (**A** and **B**), 1 μM CHI singlely or combined with 5 μM azacytidine (AZA) (**C** and **D**), AML cell lines HL60 and THP1 were collected and determined the DACT3 mRNA expression using qRT-PCR; compared with control or the combination group: ^*^P<0.05, ^**^P<0.01. (**E**-**J**) qRT-PCR analysis of the mRNA expression levels of HDAC1, 2, 3, 8 and10 in AML cell lines THP1, K562, K562/ADR, HL60, MOLM13 and MV4-11. (**K** and **L**) After treatment with different dose of CHI (0, 0.5, 2 and 5 μM) for 24 hours, the mRNA expression of HDAC1, 2, 3, 8 and 10 in AML cell lines HL60 and THP1 was determined using qRT-PCR.

To further confirm the influence of histone acetylation on DACT3 in AML, the mRNA expression levels of histone deacetylases HDAC1, HDAC2, HDAC3, HDAC8 and HDAC10 were determined in 6 AML cell lines [74-76]. As shown in Figure 6E-J, HDAC1, HDAC2 and HDAC3 are the mainly elevated deacetylases in AML, suggesting that DACT3 in AML may be mainly affected by HDAC type I. After treatment with different concentration of chidamide for 24 hours, the mRNA expressions of HDAC1, HDAC2, HDAC3, HDAC8 and HDAC10 were increased, among which HDAC1, HDAC2 and HDAC3 increased more significantly (Figure 6K and L), suggesting that HDAC type I may play the major role in regulating the expression of DACT3.

### DACT3 inhibits cell proliferation and promotes cell apoptosis of acute myeloid leukemia

In order to explore the role of DACT3 in growth of AML cells, we constructed two DACT3 overexpression AML cell lines HL60 and THP1, confirming by qRT-PCR and western blot detection (Figure S1A and B). In vitro functional experiment, it was found that the cell proliferation decreased (Figure S1A and B), apoptosis increased (Figure S1C) and cells were blocked in G0/G1 phase after DACT3 overexpression in HL60 and THP1 cells (Figure S1D). In addition, we generated cell lines derived from HL60 and THP1 that stably express a short-hairpin RNA targeting DACT3. Levels of the DACT3 mRNA and proteins were greatly diminished in two DACT3 shRNA AML cell lines compared to control shRNA cells (Figure 7A and B). Moreover, when cell growth ability was assayed, the induction of cell proliferation by adriamycin was markedly reduced in DACT3 shRNA cell lines (Figure 7C and D) and increased in DACT3 overexpression cell lines (Figure 7E and F). The above results have confirmed that knockdown of DACT3 desensitizes AML cells to adriamycin *in vitro*. To confirm this discovery, we subsequently asked whether loss of DACT3 promotes cell proliferation in AML cells *in vivo*. As shown in Figure 7G, the DACT3 shRNA HL60 tumours developed in mice are larger than the empty vector tumours (tumor weight: 0.002; tumor volume: P<0.001). Our above results suggested that, DACT3 inhibits the cell growth and promotes cell apoptosis, which is related to cell cycle arrest in G0/G1 phase.

**Figure 7.**
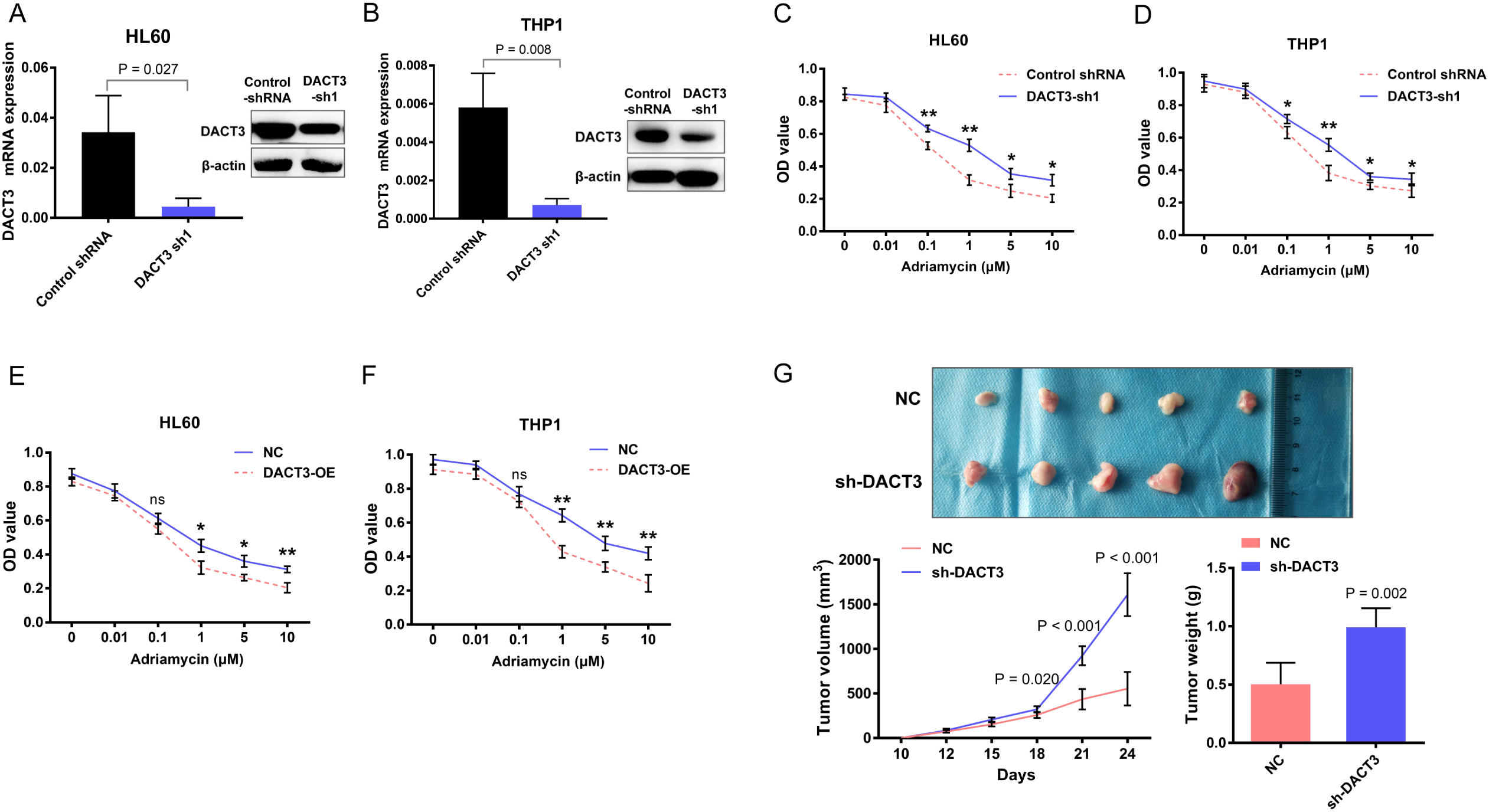
DACT3 represses the proliferation of AML cells. (**A** and **B**) After transfection with DACT3 shRNA lentiviruses, qRT-PCR and western bloting were respectively used to verify the mRNA and protein expression level of DACT3 in AML cell lines HL60 and THP1. (**C**-**F**) CCK-8 analysis of the role of DACT3 in the proliferation of AML cells: (**C** and **D**) the sensitivity of AML cells HL60 and THP1 to adriamycin after down-regulation of DACT3; (**E** and **F**) the sensitivity of AML cells HL60 and THP1 to adriamycin after overexpression of DACT3; OD value represents cell vitality. Compared with control: ^ns^P>0.05; ^*^P<0.05; ^**^P<0.01. (**G**) Down-regulation of DACT3 exhibits AML cell inhibiting in vivo: the up, comparison of tumor nodes between DACT3 knocked down HL60 cells (sh-DACT3, n=5) and HL60 cells transfected with empty vector (NC, n=5); down left, tumor volumes between sh-DACT3 and NC groups in B-NDG mice; down right, tumor weight between sh-DACT3 and NC groups.

### DACT3 is a negative regulator of Wnt/β-catenin signaling in acute myeloid leukemia cells

To explore the role of DACT3 in Wnt/β-catenin signaling in AML cells, DACT3 overexpression and DACT3 shRNA cell lines derived from HL60 and THP1 were generated as previously described. As shown in Figure 8A, compared with the control group, β-catenin protein levels were significantly decreased (HL60: P=0.002; THP1: P=0.028) in DACT3 overexpression cells, suggesting that the up-regulation of DACT3 has inhibitory effect on β-catenin. When compared with control siRNA transfecting, transfection of β-catenin siRNA into DACT3 overexpression cells evidently reduced β-catenin protein expression (HL60: P<0.001; THP1: P=0.001; Figure 8A), indicating that knockdown β-catenin enhanced the inhibition effect of DACT3 overexpression on Wnt/β-catenin signaling. Moreover, β-catenin protein level was increased (HL60: P=0.015; THP1: P=0.002) in DACT shRNA cells and decreased (HL60: P=0.003; THP1: P<0.001) when transfecting with β-catenin siRNA (Figure 8B). Further more, cell viability assay showed a significant cell growth inhibition in DACT3 overexpression cells (HL60: P<0.01; THP1: P<0.01) and β-catenin siRNA cells (HL60: P<0.001; THP1: P<0.01) compared with their control group (Figure 8C). Subsequently, we analyzed the effect of DACT3 on the Wnt/β-catenin signaling by luciferase reporter assays. As shown in Figure 8D, overexpression of DACT3 drastically suppressed Wnt signaling in HL60 and THP1 cells (P<0.01; left), whereas loss of DACT3 promoted the Wnt signaling (P<0.001; right). In summary, these results indicated that DACT3 represses Wnt/β-catenin signaling leading to inhibition of AML cell growth.

**Figure 8.**
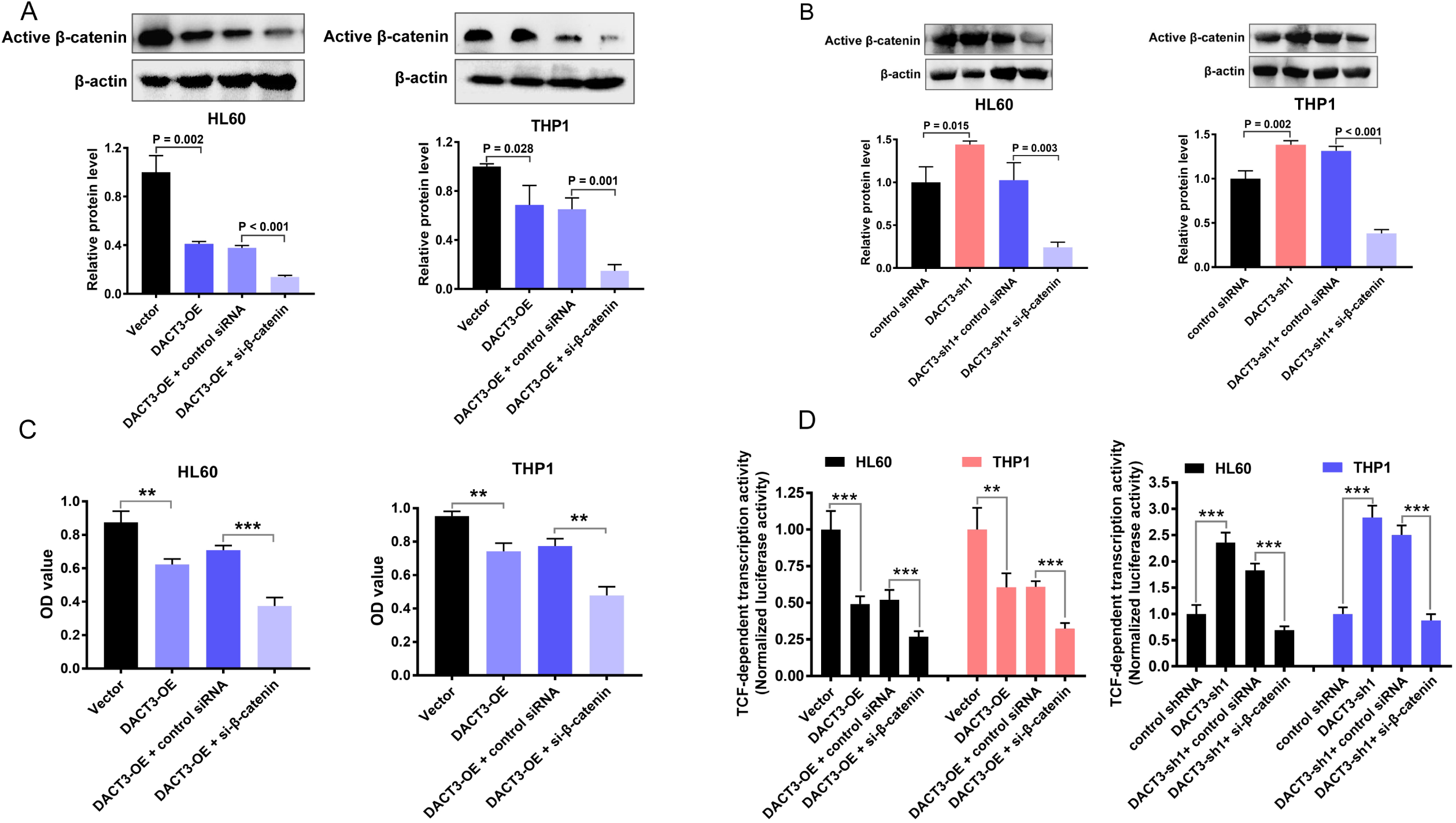
DACT3 is a Wnt/β-catenin antagonist in AML. (**A**) β-catenin protein levels in HL-60 and THP1 cells, and their DACT3 overexpression cells, DACT3 overexpression cells transfected with β-catenin siRNA or control siRNA. (**B**) β-catenin protein levels in HL-60 and THP1 cells, and their DACT3 knockdown cells, DACT3 knockdown cells transfected with β-catenin siRNA or control siRNA. (**C**) Evaluation of β-catenin knockdown effects on the growth of DACT3 overexpression AML cells HL60 and THP1 using CCK-8. (**D**) TOP/FOP dual-luciferase reporter assay analysis of Wnt/β-catenin signaling transcriptional activity in HL-60 and THP1 cells, their DACT3 overexpression cells, DACT3 knockdown cells, DACT3 overexpression or knockdown cells transfected with β-catenin siRNA or control siRNA. ^**^P<0.01; ^***^P<0.001.

### Epigenetic drugs inhibit Wnt signaling and promote apoptosis in acute myeloid leukemia cells via up regulating DACT3

Previous studies suggested that, increased DACT3 protein level can lead to degradation of Wnt signaling core component Dishevelled (DVL) protein [30]. The activation of Wnt/β-catenin signaling in AML cells can be achieved by accumulation of β-catenin in nucleus and cytoplasm [8,9]. Therefore, we studied the effect of epigenetic drugs treatment on DACT3 and β-catenin level. Consistent with the previously observed changes in transcriptional levels (Figure 6C and D), western blotting analysis confirmed that DACT3 protein levels in HL60 and THP1 cells significantly increased after co-treatment with chidamide and azacytidine (Figure 9A). On the contrary, immunofluorescence analysis showed that the co-treatment resulted in a significant decrease of intracellular total β-catenin expression in HL60 and THP1 cells (Figure 9C), suggesting that the co-treatment of chidamide and azacytidine inhibits Wnt/β-catenin signaling in AML. In addition, the co-treatment of the two epigenetic drugs synergistically increase AML cell apoptosis (Figure 9B). Further more, we inquired the relationship between the expression of DACT3 and other core components of Wnt signaling in AML patients. As shown in Figure 9D, the low expression of DACT3 in AML patients was usually accompanied with the up-regulation of active β-catenin (non phosphorylation), DVL2, and the downstream target LEF1 of Wnt signaling, which have been confirmed in our previous study [46].

**Figure 9.**
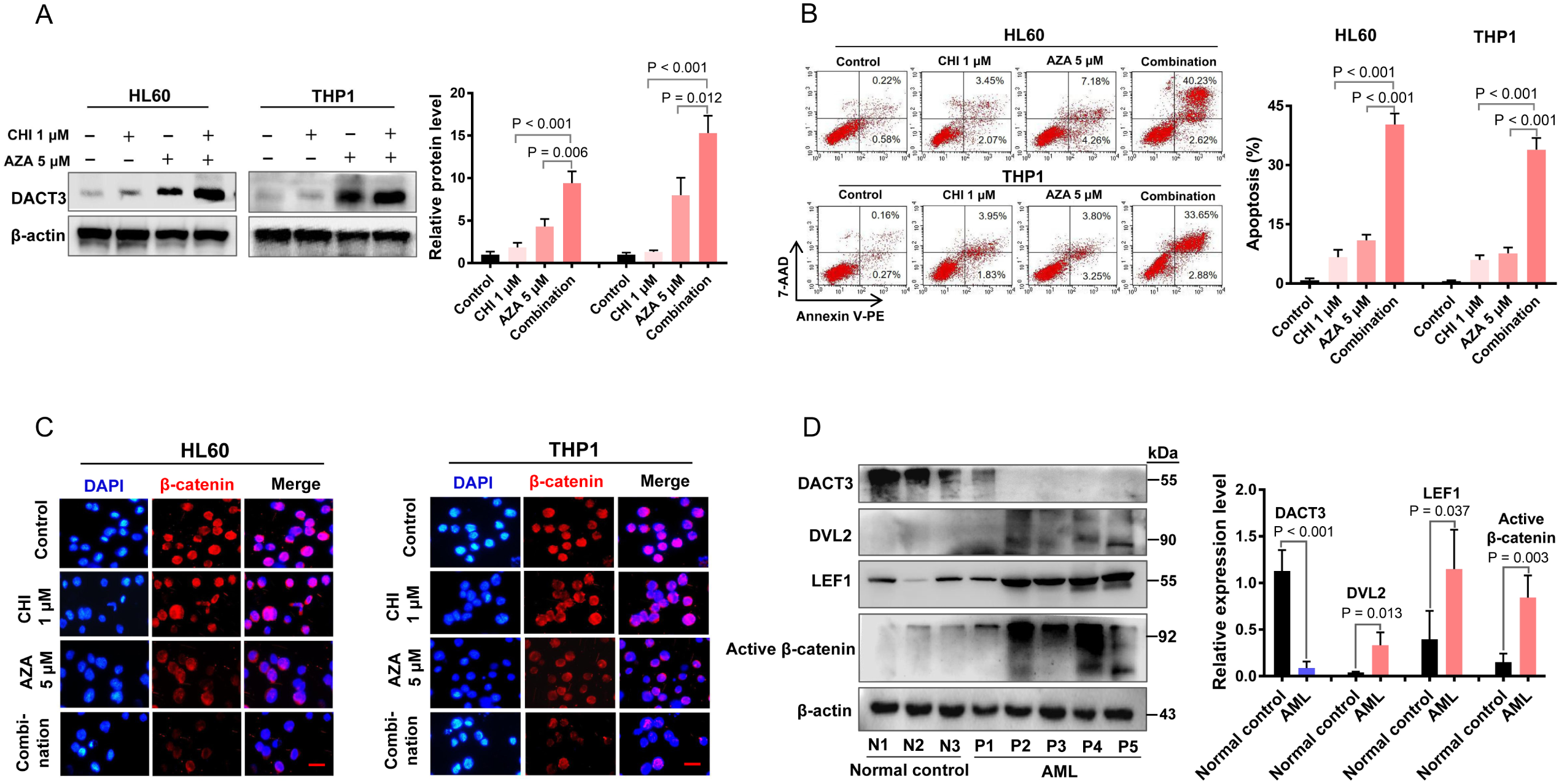
Epigenetic drugs inhibit Wnt signaling and promote apoptosis of AML cells by up regulating DACT3. After treatment of HL60 and THP1 cells with 1 μM chidamide (CHI) singlely or combined with 5 μM azacytidine (AZA) for 48 hours, (**A**) the DACT3 protein levels were evaluated with western blotting, (**B**) cell apoptosis was assessed using flow cytometry, (**C**) β-catenin expression level was determined using immunofluorescence analysis. (**D**) Protein levels of Wnt signaling components DACT3, DVL2, LEF1 and active β-catenin (non phosphorylation) in AML patients (n=5) and normal controls (n=3) were analyzed by western blotting.

### Dual epigenetic treatment represents a promising therapeutic strategy for the treatment of acute myeloid leukemia with FLT3 mutations

In line with the above discoveries, we found that co-treatment with chidamide and azacytidine led to greater reduction in tumour growth than monotherapy (Figure 10A-C), indicating a combined anti-leukemia activity of the two epigenetic drugs *in vivo*. Next, we collected five relapse/refractory AML patients with FLT3-ITD mutation (Supplementary Table S3). Notably, we have successfully applied the double epigenetic drugs based salvage chemotherapy in treating relapse/refractory patients from Changsha and Hainan, China, and some of them obtained complete remission (Figure 10B, unpublished data). Together with previous results, our data suggest that the dual epigenetic treatment with chidamide and azacytidine is a promising therapeutic strategy for the treatment of AML with FLT3 mutations.

**Figure 10.**
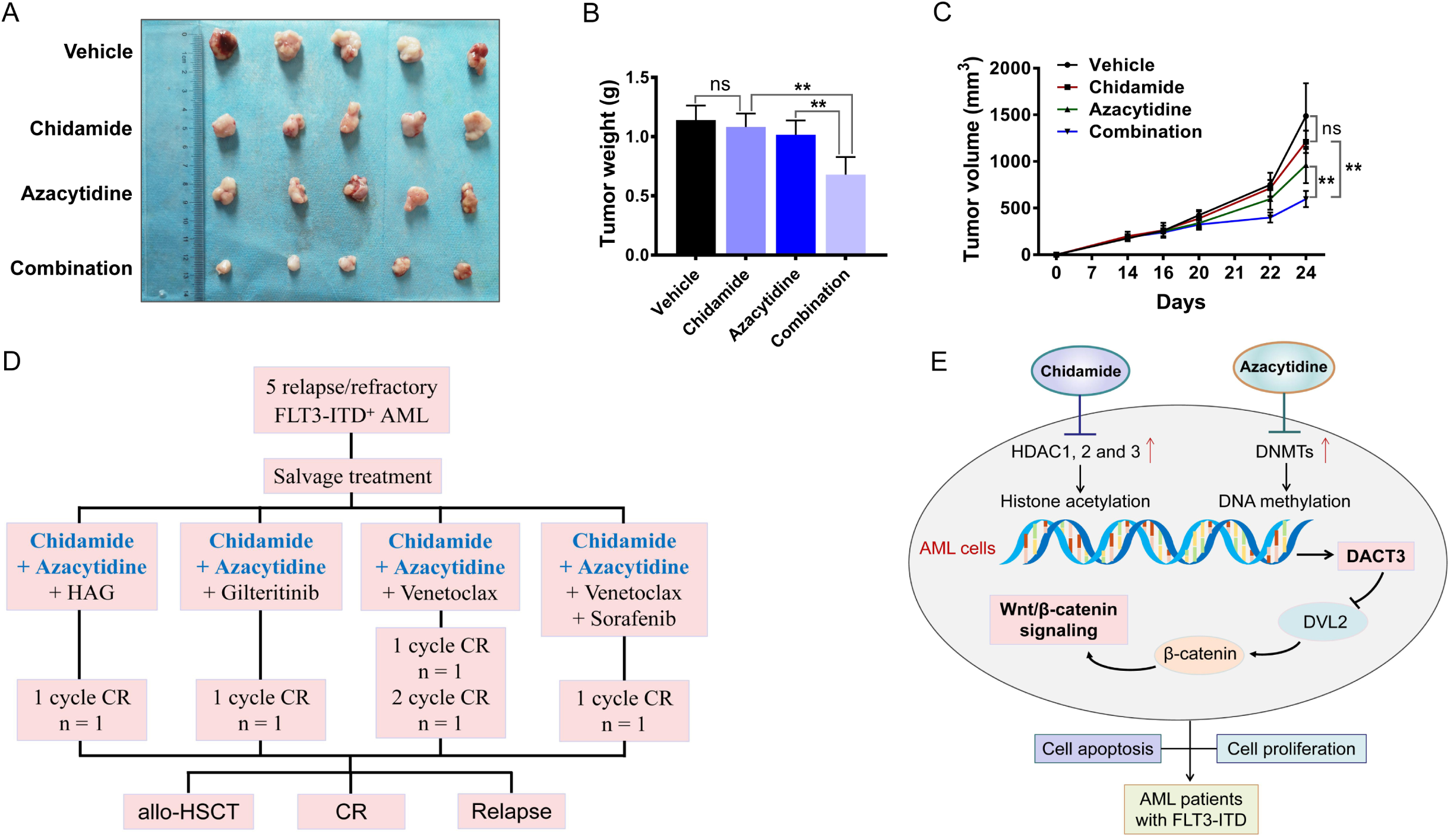
Double epigenetic treatment leads to Considerable therapeutic response of FLT3-ITD AML. (**A**) Chidamide and azacytidine exhibit synergistic anti-leukeia effect in vivo. Tumor volumes (**B**) and tumor weight (**C**) of HL60 knockdown cells following treatment with single drugs and combined drugs (n=5). ^ns^P>0.05; ^**^P<0.01. (**D**) The double epigenetic drugs based treatment regimen used for five relapsed/refractory FLT3-ITD AML patients in clinic. HAG: homoharringtonine plus cytarabine and granulocyte stimulating factor; CR: complete remission; allo-HSCT: allogeneic hematopoietic stem cell transplantation. (**E**) The model of our working hypothesis.

## Discussion

The aberration of Wnt/β-catenin signaling has been detected in leukemia, myeloma and lymphoma [77], and the β-catenin is highly expressed in most AML cells, especially in bone marrow leukemic stem cells (LSC) [15]. In the present study, we found that the Wnt/β-catenin signaling is active in AML cells. Importantly, we also demonstrated that DACT3, an antagonist of β-catenin, [10,30,78], inhibits cell growth and promotes apoptosis via inhibiting Wnt/β-catenin signaling in AML cells. The combination of DNA methyltransferase inhibitor (DNMTi) and histone deacetylase inhibitor (HDACi) cause upregulation of DACT3 resulting in anti-leukemia effect (Figure 10E).

Previous studies have found that, the expression of DACT1 or DACT2 was reduced in various solid tumors, and was identified as a tumor suppressor genes [26,27,79]. It has been reported that DACT3 is transcriptionally inhibited and is a negative regulator of the Wnt/β-catenin signaling pathway in colorectal cancer [30]. Recently, Gao et al. found that DACT3 is low expression in AML and negatively adjusts Wnt/β-catenin pathway via downregulation of DVL2 in AML. However, the role of DACT3 is not fully understood in AML for their limited samples and design. For this study, we first inquired differential expression of DACT genes in AML patients by RNA-seq of a small sample, and hierarchical clustering analysis showed that DACT3 was the most differentially expressed one among the three homologous genes. Then qRT-PCR analysis of a expanded AML cohort validated that the DACT3, other than DACT1 or DACT2, was the differential and down-regulated DACT gene in AML, indicating that DACT3 plays the crucial role. The online differential expression analysis based on TCGA database, DACT3 mRNA and protein expression analysis in 6 AML cell lines, 129 newly diagnosed and 13 relapsed AML patients further supports this conclusion. In addition, we also observed that DACT3 was significantly reduced accompanied by up regulation of components representing Wnt signaling activation [Figure 1A and Figure 9D], which is consistent with prior studies [25]. It is worth noting that, our data showed that DACT3 expression decreased after complete remission and increased again at relapse, suggesting that DACT3 level may be related to therapeutic effect of AML. More importantly, DACT3 level was associated with drug resistant in AML. Moreover, we observed that low DACT3 expression was associated with higher white blood cell count and BM blasts, younger age, FLT3 and CEBPA gene mutation in AML. Survival analysis revealed that patients with low DACT3 expression had a poor prognosis in non-M3 AML, whereas DACT3^low^ had a shorter OS in AML cases from Genomoscape dataset.

Wnt signalling is necessary for the maintenance of leukaemic stem cells [12]. Promoter methylation of genes in Wnt signaling (e.g., sFRP1, DKK1, DKK3, SOX17, WIF1, cyclin D1, LEF1) causes genes to be down regulated in their mRNA expression, thereby deregulating the Wnt signaling [35]. In this study, we found that the expression of Wnt inhibitors SFRP1 and DKK1 decreased in AML patients, while the expression levels of Wnt upstream signal molecules WNT6, WNT7B, FZD2, DVL2, PRICKLE1 [46] and Wnt downstream effectors CCND1, MYC and CD44 increased (Figure 1A). These results indicate that Wnt/β-catenin signaling may be activated in AML patients. Prior studies have found that DACT3 inhibits TCF dependent transcription through interaction with DVL2 in cancer [30]. Through *in vitro* function experiments and β-catenin interference experiments, our data showed that DACT3 inhibits cell proliferation and promotes cell apoptosis, and acts as a negative regulator of Wnt/β-catenin signaling in AML cells. In addition, the inhibition of DACT3 on the growth of AML was further confirmed in the subcutaneous xenograft tumor model of B-NDG mice. Our study suggests that DACT3 may be a potential target for AML treatment.

Epigenetics is known to have three regulatory forms, histone methylation or acetylation modification, DNA methylation and chromatin remodeling [80-82]. Based on the previous relationship between mRNA and protein expression of DACT3 (Figure 2H), we explored the regulatory mechanism of DACT3 transcriptional inhibition in AML from two aspects: DNA methylation and histone acetylation. Validation from RQ-MSP analysis of 85 de novo AML patients and qRT-PCR analysis of AML cell lines treated by chidamide and azacytidine, our results showed that loss of DACT3 in AML is associated with DNA methylation, and DACT3 expression is affected by histone acetylation. Previous studies analyzed genetic and epigenetic changes of Wnt signaling genes and DNA hypermethylation was found to be enriched in AML [13]. Our data showed that azacitidine, a therapeutic agent with DNA hypomethylating activity[83,84], can potently up regulate the DACT3 and inhibit β-catenin leading to apoptosis increasing in AML cells. Further investigations were warranted to confirm that chidamide, a novel HDACi targeting HDAC1, 2, 3, and 10 [85], was able to enhance the above effects of azacitidine. These results suggest that epigenetic drugs inhibit Wnt signaling and promote apoptosis in AML cells via up regulating DACT3.

Wnt activation occurs as the result of mutations in AML, and mechanisms have been described such as Wnt antagonist inactivation through promoter methylation [12]. AML with mutations of FLT3-ITD, c-Kit, TP53, etc. often has poor response to chemotherapy, high relapse rate and aggressive progression [1]. Disruption of Wnt/β-catenin exerts anti-leukemia activity and has synergistic effect with FLT3 inhibition in FLT3-mutant AML [86-88]. Current AML treatment protocols rely on intensive chemotherapy therapies [89]. Therefore, we have tried to apply the double epigenetic drugs (chidamide and azacitidine) based salvage chemotherapy in treating relapse/refractory patients, and obtained a promising therapeutic response (Figure 10D).

In summary, our results indicated that DACT3 is significantly down regulated in AML and low DACT3 level is related to poor survival. DACT3 plays a negative role in regulating Wnt signaling and cell growth in AML. In addition, the co-treatment of chidamide and azacytidine up-regulates DACT3 expression and promotes AML cell apoptosis via inhibiting Wnt/β-catenin signaling in AML cells, indicating a promising therapeutic prospects in FLT3-mutant AML. More importantly, our studies suggested that DACT3 is a candidate gene for therapeutic target and targeting DACT3 has the potential to validate the combination therapy of DNMT inhibitors and HDAC inhibitors.

## Acknowledgments

We thank Yinghong Zhu for the retrovirus constructs, Suo Liu and Qiangqiang Zhao for help with mouse model constructs, Hui Zeng for the AML cell lines, Yanjuan He for helpful discussion, Xiaotao Wang for his initial insight, and Xin Li for technical assistance.

## Authors’ contributions

DFJ and QYM performed the experiments and wrote the manuscript. HGS, JM, RLZ, XXJ, CJL and WYL provided technical support for this study and participated in discussions about this study. PFC and MD designed the research and edited this manuscript. All authors read and approved the final manuscript.

## Ethics approval and consent to participate

The study was approved by the Institutional Ethics Committee of the Third Xiangya Hospital of Central South University. Informed consent from patients or their legal guardians were obtained according to the Declaration of Helsinki. Animal experiments were approved by the Institutional Animal Care and Use Committee of Central South University.

